# Action history and target uncertainty co-determine human reaching direction under time pressure

**DOI:** 10.1101/2023.12.17.572036

**Authors:** B Keane, L Smith, TJ Carroll

## Abstract

Given the inaccuracies that are inherent in biological sensory and motor functions, animal sensorimotor control should be probabilistic. Effective neural control systems for movement must estimate the most likely true state of the world on the basis of uncertain sensory information, and to select and execute movements that are most likely to be successful given motor variability. Bayesian inference dictates that if sensory information is ambiguous (e.g. low light conditions), animals should rely more on their past experience of target locations to guide motor planning, and less on their current sensory information about target location. Here we investigated how time pressure affects degree to which the precision of sensory information about a target influences movement direction bias towards previous target locations. We used a paradigm developed by Dekleva et al. (2016) that involved uncertain cues to the location of a hidden target for reaching with movement preparation time strictly controlled. One group of participants (n=10) were required to initiate their reaches within 150-300ms of target presentation, and a second group (n=10) were required to initiate their reaches with 1150-1300ms of target presentation. We found that participants relied more on prior target location information when target precision was reduced under time pressure, which suggests that integration of target uncertainty information according to Bayesian principles is an inherent component of sensorimotor transformation and does not require time-consuming cognitive processes.

## Introduction

Animal survival depends upon the ability to execute movements that are customized to the current environmental context. Decisions about what type of movement to make should depend, for example, upon whether there is a predator or food source nearby. The details of selected movements should take into account the identity, location and motion of physical stimuli in the animal’s vicinity, as well as the current physical state of the animal itself. At best, these requirements present a complicated problem for neural systems; to link environmental and internal states with specific actions. However, information about the external world and the body can only be obtained through the senses available to an organism, and all sensory systems provide only imperfect estimates of reality. Even further, the processes of motor execution and planning are inherently noisy, so decisions about what to do will not be perfectly implemented. The neural processes of movement selection and control should therefore be probabilistic to maximize the chances of organism survival and species continuation. That is, a key function for neural systems such as the vertebrate brain should be to estimate the most likely true state of the world on the basis of uncertain sensory information, and to select and execute movements that are most likely to be successful given motor variability.

The framework of Bayesian inference defines how animals should make statistically optimal decisions given uncertain information. Bayesian principles illustrate how the most accurate estimate of the state of the environment takes into account not only current sensory signals, but also the probability that different environmental states will occur. For example, if we consider the task of judging the true location of an object in the visual field *θ*, on the basis of sensory signal *x*, Bayes’ rule (eq. 1, Bayes and Price, 1763) states that the probability that an estimated location is correct, given the sensory data (known as the *posterior probability*), is proportional to the probability that the sensory data provides accurate evidence about the true location (known as the *likelihood*) multiplied by the probability of the true location before the sensory data were observed (known as the *prior probability*).

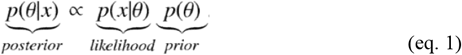

This implies that if an animal is to identify the best possible estimate of the true object location (or the *maximum a posteriori* estimate of *θ*), then it should weight the two sources of evidence for *θ*; the sensory signal *x*, and the prior probability of *θ*, by their respective precisions. Thus, if sensory information is ambiguous (e.g. low light conditions) then the animal should rely more on its past experience of visual scenes to estimate object location, and less on the visual signals it receives.Conversely, uncorrupted visual information, or a broad distribution of probable object locations, should shift the animal’s estimate toward its current sensory data at the expense of its past experiences.

Given the competitive pressures of natural selection, it is perhaps not surprising that animals show many hallmarks of Bayesian inference in their sensorimotor behaviour. For example, when humans make online corrections to reach trajectories on the basis of uncertain visual feedback of their hand location, their judgements are intermediate between the visual stimulus location and the average of the prior distribution of endpoint locations (Kording and Wolpert, 2004). Crucially, the degree to which spatial estimates are biased toward the prior distribution versus the current sensory information depends on the quality of the visual stimulus; blurry, unreliable stimuli were judged closer to the prior than clear, unambiguous stimuli. Tassinari et al (2006) also found that the degree to which humans biased the endpoint locations of their pointing movements on the basis of a prior distribution of targets that was explicitly displayed to them was proportional to the precision of the partial information provided about target location. Thus, it appears the humans appropriately account for the likelihood of sensory information about movement targets and feedback.

Human reaching is also more accurate toward visual targets in the vicinity of previously executed movements, whereas reaches toward rarely experienced targets are biased towards the location of past (Verstynen and Sabes, 2011). Similarly, human judgements of orientation are biased toward the Cardinal axes, where the highest proportion of line orientations are evident in natural scenes (Girshick et al., 2011). The behaviour and neural activity of many other animals conform to Bayesian principles, including biases in owl sound localization toward central directions in space (Fischer and Pena, 2011), and convergence of spontaneous visual cortex activity patterns toward patterns evoked by natural scenes in developing ferrets (Berkes et al., 2011). Thus, there is evidence that animals take account of both prior and likelihood components of evidence to guide sensorimotor performance. There is also extensive evidence that animals optimally integrate information from multiple sources of sensory information (e.g. visual and somatosensory information about target location) according to their relative precisions according to Bayesian statistical principles (van Beers et al., 2002, Ernst and Banks, 2002, Fetsch et al., 2011, Jacobs, 1999).

One question that remains from studies in which animals have ample time to consider their aiming direction before moving, is the extent to which Bayesian inference is the product of cognitive reasoning versus low-level sensorimotor transformation. The work of Dekleva et al. (2016) is relevant to this issue, as they had monkeys move toward the location of hidden targets while systematically manipulating the visual uncertainty of the target location by showing scattered cues on a screen. These cues provided only an approximation of the hidden target’s location as they were randomly drawn from a distribution centered on the target location. Once the reach was performed, reward was provided contingent on the correspondence between the actual reach angle and the hidden target location. In many experimental sessions, the monkeys showed behaviour consistent with the predictions of Bayesian integration of sensory information and prior target location; that is a greater bias toward the prior when likelihood information was less precise. However, there were also some experimental sessions in which the monkeys displayed idiosyncratic behaviour that did not take into account the target likelihood for reach direction. As the animals had 700-1000ms to view target likelihood before they made a response, it seems possible that the idiosyncratic behaviour could reflect variations in cognitive approaches to the task.

One way to infer the degree to which cognitive factors influence motor behaviour is to restrict the time available for sensorimotor processing of the target. Marinovic et al. (2017) and Reuter et al. (2019) used this approach to confirm that findings of Verstynen and Sabes (2011) that reaching is biased toward the center of the the prior distribution of targets when there is pressure to respond quickly to the target presentation. By contrast, reaching biases and saccade direction biases were dramatically reduced when participants had ample time to view the target before moving. Thus, the degree to which humans apply Bayesian inference appears subject to the availably of time to prepare movements after a target is presented. More specifically, priors seem to influence movement execution through a combination of temporally-stable processes that are strictly use-dependent, and time-dependent processes that reflect prediction of future actions.

In the current experiment we aimed to investigate how time pressure affects degree to which the precision of sensory information about a target (i.e. the likelihood information) influences movement direction bias towards previous target locations (i.e. the prior information). We used the paradigm reported by Dekleva et al. (2016) involving multiple target cue markers draw from a Gaussian distribution centered on the true target location, and had participants reach with movement preparation time strictly controlled. We predicted that participants would rely more on prior target location information when target precision is reduced under time pressure, which would imply that integration of likelihood information is an inherent component of sensorimotor transformation and does not require time-consuming cognitive processes.

## Methods

We recruited 20 right-handed participants (15 female, 18-32 years of age) who were naïve to the purposes of the experiment from the University of Queensland undergraduate Psychology student cohort. Participants provided written informed consent prior to any data collection procedures. Testing sessions took approximately hour to complete, and participants received course credit for their time.Participants were randomly allocated to one of two between-subject conditions (preparation time: short or long, 10 participants each) prior to testing. This study was approved by the University of Queensland Human Research Ethics committee.

### Motion-capture procedure

We first attached a small motion-capture marker to the tip of participants’ right index. This marker allowed us to track the location of the participant’s hand in a pre-calibrated space, actively monitored by a set of OptiTrack Flex cameras. We used OptiTrack’s Motive (v1.8) software to reconstruct the calibrated space and compute marker-coordinates, which we then broadcast to MATLAB using the OptiTrack NatNet SDK.

Participants were asked to sit at a comfortable height so that they could look down into a semitransparent mirror, reflecting a computer monitor suspended above (see Figure 1, below). When each participant first sat down the mirror was not occluded, meaning participants could see their hand through the glass, as well as a faint reflection of the image on the monitor. We first conducted a calibration procedure with each participant, to map the motion-capture coordinates to screen-coordinates. Participants were presented with a grid of nine small white circles in series, and were asked to align the marker on their finger with the circle visible in the reflection of the monitor while keeping their finger touching the table surface. We then linearly mapped the motion-capture coordinates from each of these nine locations to the screen-coordinates of the circles, and added an occluder to the back of the mirror that remained in place for the duration of the testing session.

**Figure 1.**
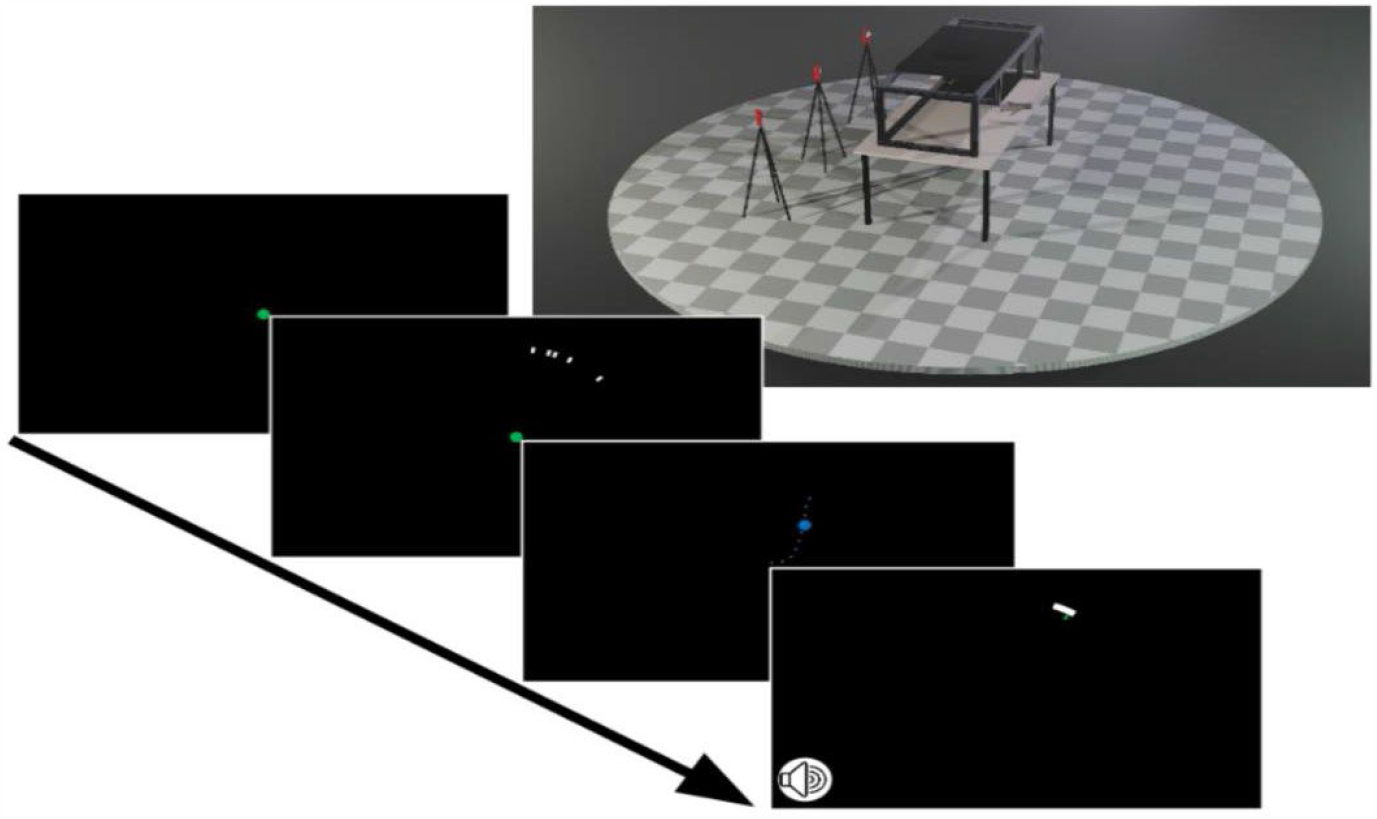
Stylised render of the testing setup and trial progression. Note that the mirror used was semitransparent; when not occluded participants could see their hand in the tracking space, allowing us to conduct calibration procedures. The inserts illustrate the progression of information provided to participants during each trial. They first had to return their hand to a central home marker, before 5 target markers illustrated the approximate location of a hidden target. They were given feedback of their hand position during movement (blue dot), and of the final position of their hand with respect to the target at the required movement extent.

To validate the calibration measurements, we then repeated the calibration process with the hand occluded by the mirror, this time presenting a red circle to indicate the calibrated marker location. This procedure provided veridical two-dimension visual feedback of marker location while the participants’ finger remained on the elevation plane of the reflection of the monitor. We stressed that participants must keep their finger touching the table during their reaching movements, as this approximately matched the elevation of the marker with the elevation of the reflected monitor, meaning visual feedback would be (very close to) veridical.

### Experimental Procedure

Participants performed a total of 360 reaches towards hidden targets that were approximately indicated by the location of five cue markers spaced on an arc at the required reach distance. The locations of the cue markers were randomly drawn from a Gaussian distribution centred on the location of the hidden target. The precision of the distribution was manipulated within-subjects at three levels: standard-deviations (SD) of 5, 30, and 50 degrees. The participants were informed that the five cue markers were not the target, but that they provided some information as to where a hidden target might be. After making each reaching movement, the actual location of the hidden target was revealed, and participants were able to compare their reach direction at the required target distance with the target location, as shown in Figure 1. Participants were required to commence their reaching movement no earlier than 50ms and no later than 100ms from a start tone. The start tone was preceded by three preparatory tones, each spaced 500ms apart, to allow the participant to anticipate the start tone. Participants were split into a short preparation (200ms, n=10) or long preparation (1200ms, n=10) group. The available times from target presentation to movement initiation therefore varied between 150-300ms in the short preparation task, and 1150-1300ms in the long preparation task.

### Data Analysis

The primary outcome measure was the reach angle between the target and the point at which the finger crossed the arc defining the required movement extent. We obtained the slope of a linear model relating the centroid of the visual cues on context trials to participants’ reach angle would vary as a function of visual uncertainty and preparation time. We used least-squares regression to fit a linear model with a slope and intercept term to these values for each participant, independently for each level of visual uncertainty. We also fit this model to data measured on probe trials. This approach is similar to that of Dekleva et al. (2016). In this case, a slope of 1 indicates participants reached directly toward the centroid of the visual cues (perhaps with some uniform global bias, indicated by the intercept term), with no bias toward prior target locations. A slope of zero, by contrast, indicates participants’ reaches were, on average, directly toward the prior target location (again, allowing for a global rotational bias, captured by the intercept term). We tested whether the slope differed between target uncertainty levels and movement preparation time conditions with a two-way mixed-effects analyses of variance (ANOVA). We use the same two-way ANOVA model to check if the standard deviation of reach angles differed between target uncertainty levels and movement preparation time conditions. Statistical analyses were conducted using JASP. Alpha was set to 0.05, and the Greenhouse-Geisser correction was applied to the degrees of freedom when the assumption of sphericity was violated.

## Results

The degree of target uncertainty was expected to affect the variation in reach directions. We tested whether this effect was different for the long and short preparation groups with a three (likelihood SD: 5, 30, 50) x two (preparation time: short, long) mixed-effects ANOVA on the standard deviation of reach direction angles (see Fig 2 for long preparation group, Fig 3 for short preparation group data, and Fig 6 for a direct comparison between groups). There was a significant main effect for target uncertainty (*F*1.6, 27.9 = 342.2, *p* < 0.001), but no main effect of preparation time (*F*1, 18 = 12.2, *p* = 0.97), or interaction effect (*F*1.6, 27.9 = 0.52, *p* = 0.56). Thus, there was a clear trend toward increased reach direction standard deviation as the target uncertainty increased for both movement preparation time groups, as expected. Interestingly, the availability of time to process the target features before movement initiation had no discernible effect on movement precision.

**Figure 2.**
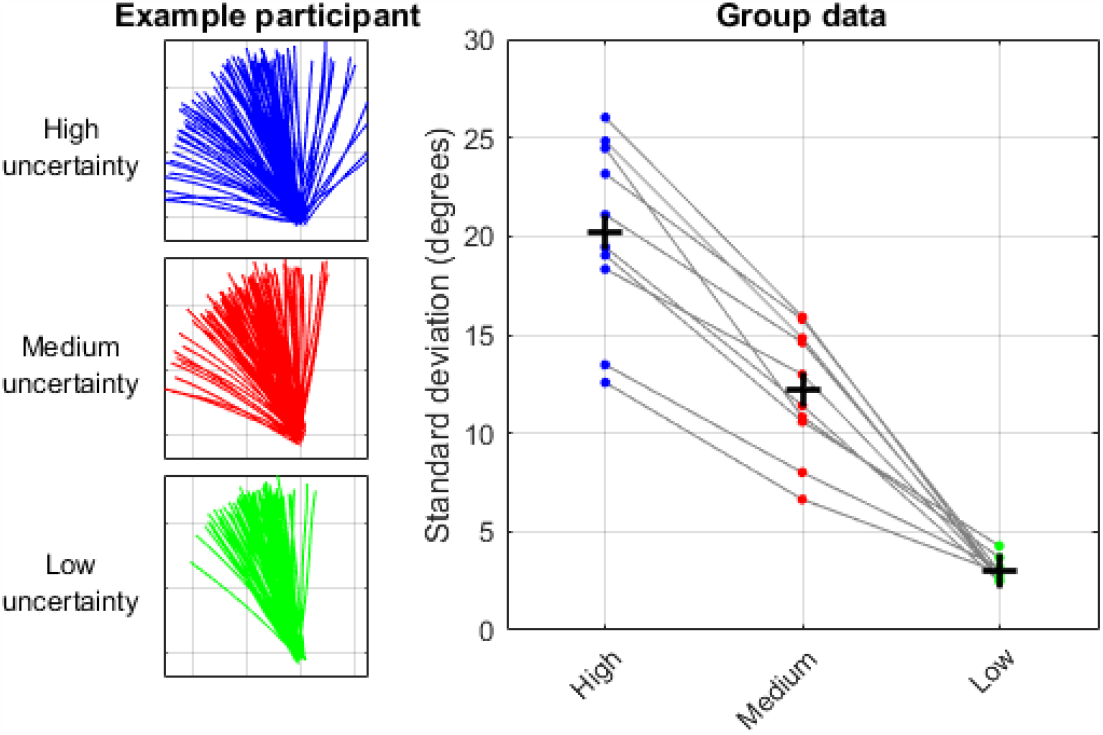
Left panels show each trajectories for an example participant in the long preparation condition for the three levels of target uncertainty. The right panel shows the standard deviation of reach directions for each participant at the different uncertainty levels (small coloured dots) and the group mean SDs (large crosses).

**Figure 3.**
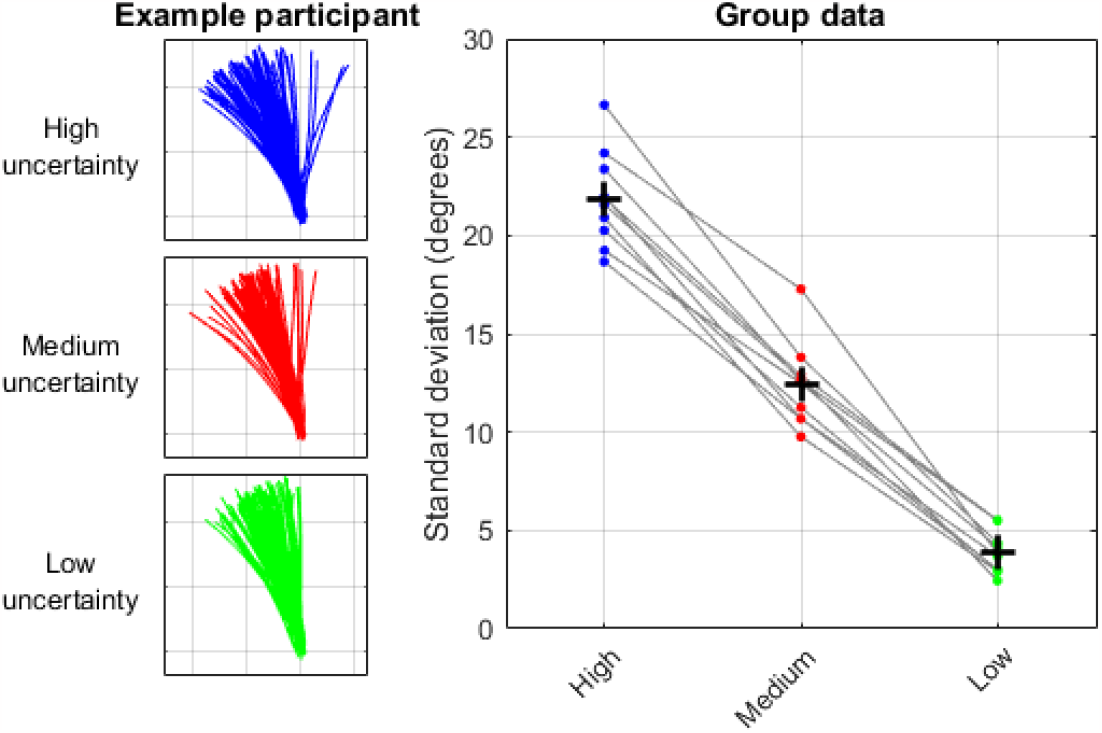
Left panels show each trajectories for an example participant in the short preparation condition for the three levels of target uncertainty. The right panel shows the standard deviation of reach directions for each participant at the different uncertainty levels (small coloured dots) and the group mean SDs (large crosses).

The primary measure of the degree to which participants took account of the target uncertainty according to the principles of Bayesian inference is the slope of linear relationship between the angle of the cue centroid and the reach direction angle (see figure 4 for the long preparation group, figure 5 for the short preparation group, and figure 6 for a direct comparison between groups). A slope of 1 indicates participants reached directly toward the centroid without an influence of prior target locations, and a slope of zero indicates that participants reached directly toward the prior target location. The three (likelihood SD: 5, 30, 50) x two (preparation time: short, long) mixed effects ANOVA showed significant main effects of cue marker SD (F1.51, 27.19= 36.0, p < 0.001) and preparation time group (F1, 18 = 12.2, p = 0.003). Thus, as previously reported, reach directions were more biased toward the prior for uncertain targets (Dekleva et al., 2016, Kording and Wolpert, 2004, van Beers et al., 2002, Ernst and Banks, 2002, Tassinari et al., 2006), and bias towards the prior was greater when time available for sensorimotor transformation was short (Marinovic et al., 2017, Reuter et al., 2019). However, there was also a significant interaction between the SD condition and preparation time (F1.51, 27.19 = 7.0, p = 0.007), indicating that likelihood information was not integrated in the same way for the two preparation time groups. The effect of target uncertainty on slope was far more consistent across participants in the short preparation group, with all but one of the ten showing a monotonic reduction in slope with decreasing target uncertainty. By contrast, the effects of target uncertainty were highly variable for the long preparation group, with some participants showing little effect of target uncertainty.

**Figure 4.**
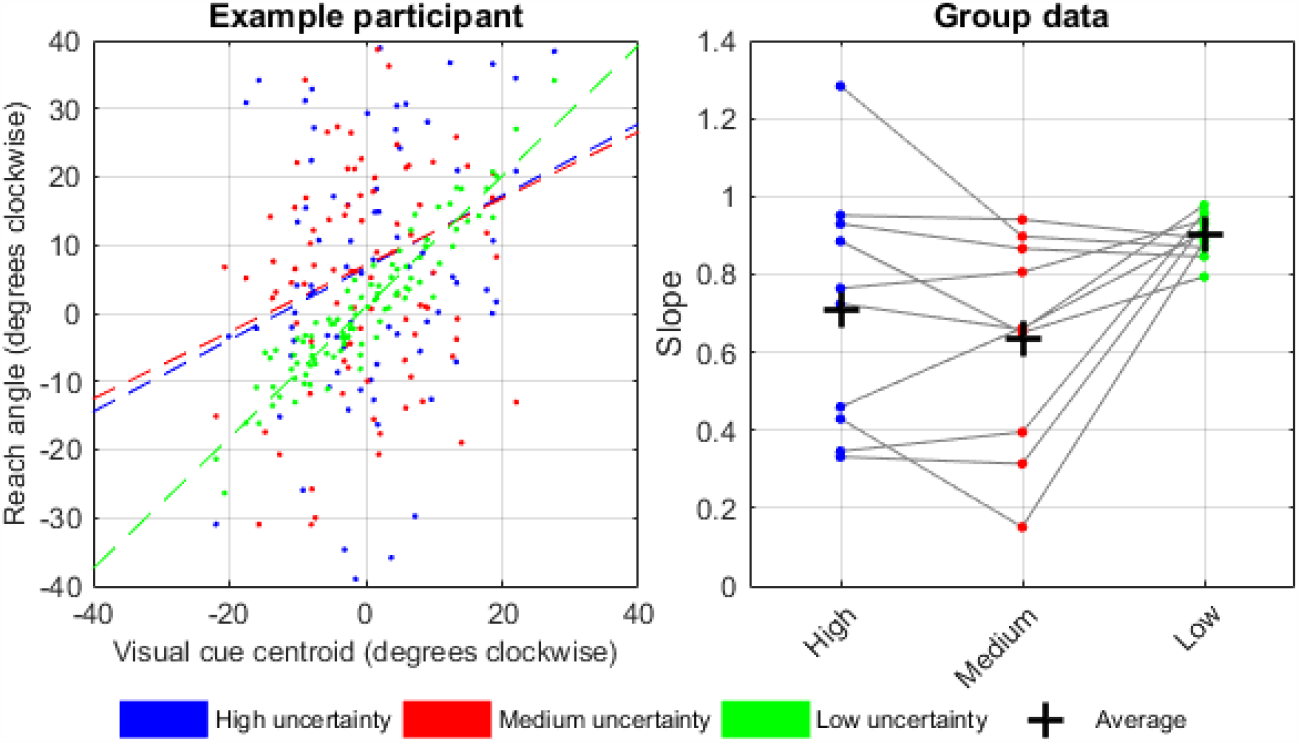
Left panel shows the scatterplot of reach direction versus visual cue centroid direction for an example participant in the long preparation condition for the three levels of target uncertainty. The right panel shows the slope of the relationship for each participant at the different uncertainty levels (small coloured dots) and the group mean SDs (large crosses).

**Figure 5.**
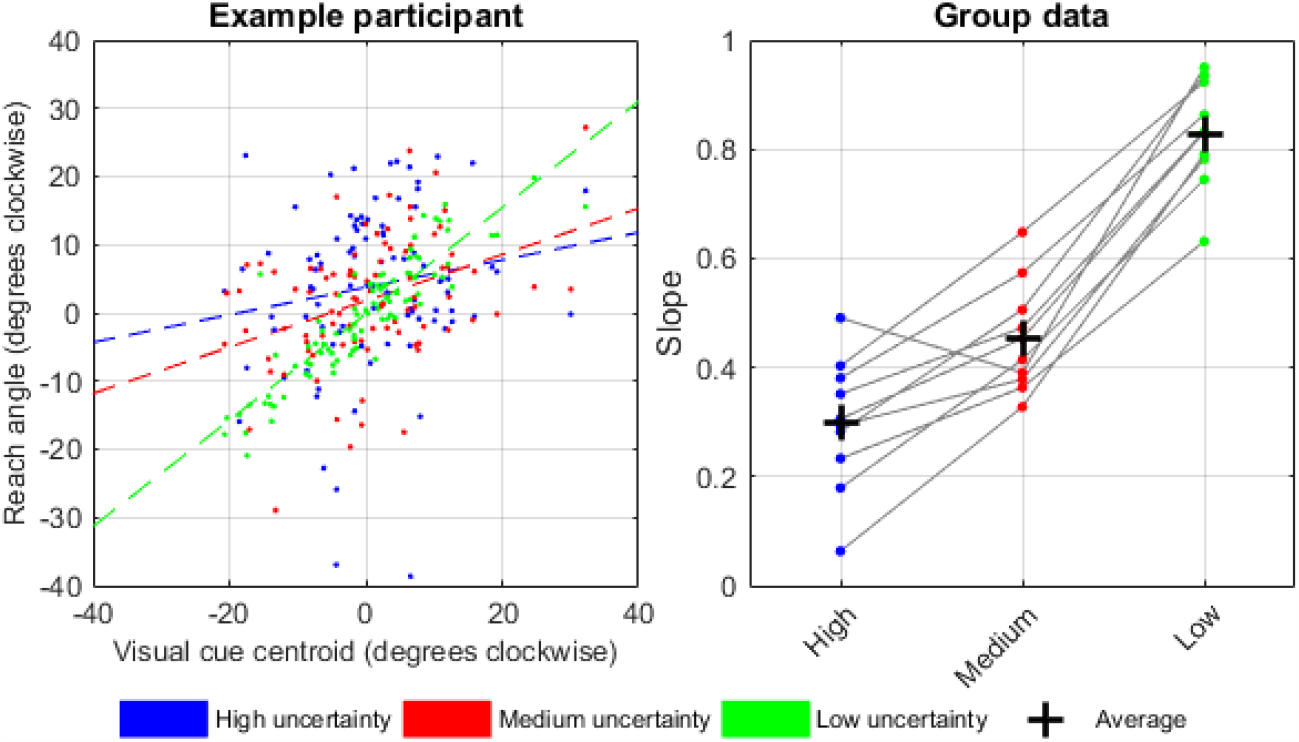
Left panel shows the scatterplot of reach direction versus visual cue centroid direction for an example participant in the short preparation condition for the three levels of target uncertainty. The right panel shows the slope of the relationship for each participant at the different uncertainty levels (small coloured dots) and the group mean SDs (large crosses).

**Figure 6.**
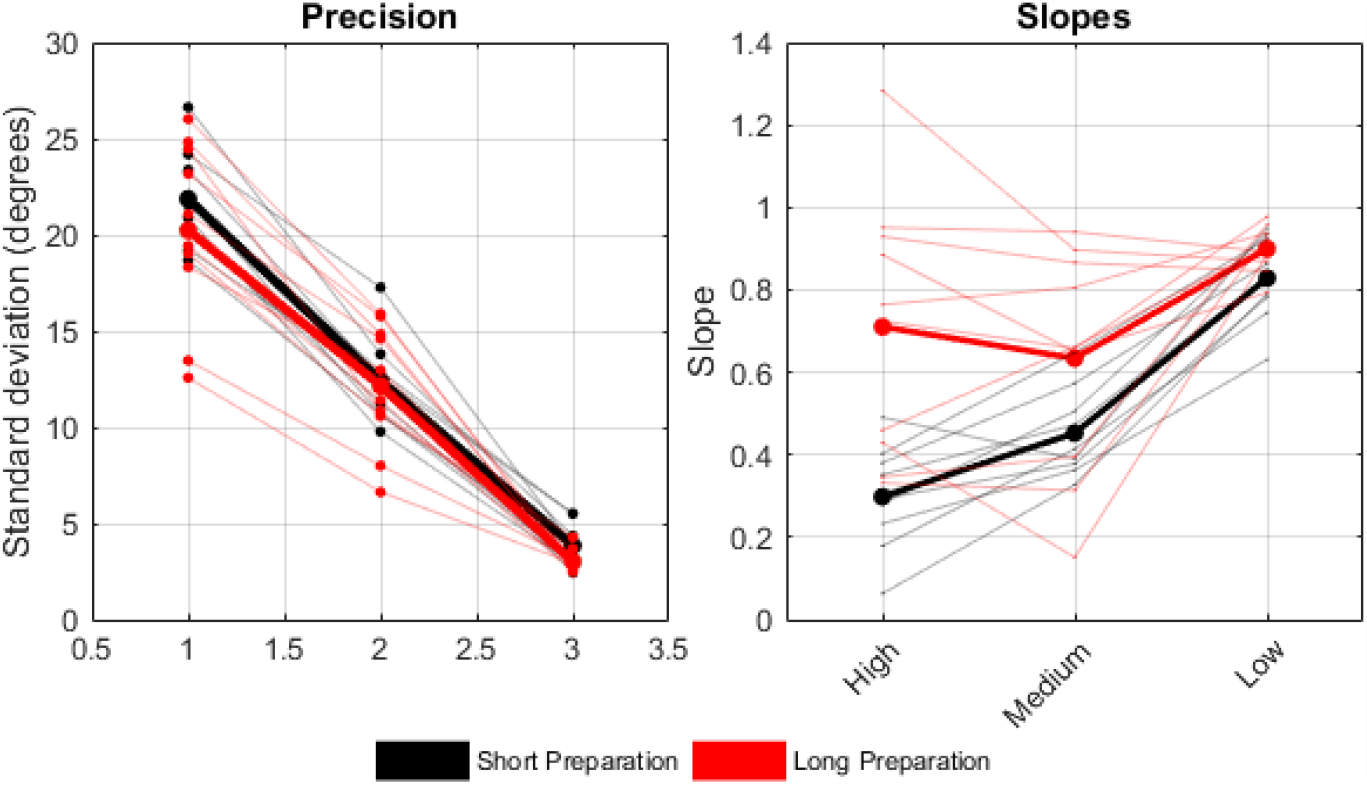
Comparison plots showing reach standard deviations (left plot) and slopes of the reach direction versus cue centroid direction linear fits (right plot) for the short and long preparation groups.

## Discussion

The key result of the current study is that humans weight prior and likelihood information in qualitative agreement with Bayesian inference when there is little time available for cognitive processing of target features before movement initiation. This suggests that Bayesian inference can be the product of lowlevel sensorimotor transformation processes and does not require time-consuming cognitive reasoning. Indeed, the effect of target uncertainty on reach bias was highly variable across participants when we enforced a long movement preparation time, suggesting that cognitive processes might lead to idiosyncratic strategies to integrate target uncertainty and action history. This interpretation might help explain the results of Dekleva et al. (2016), who showed that monkeys behaved in a manner inconsistent with Bayesian inference in some experimental sessions.

Previous work involving control of sensorimotor preparation time showed that there are two distinct neural processes that contribute to reaching bias toward the prior distribution of targets (Marinovic et al., 2017, Reuter et al., 2019). It is important to note that in these experiments, targets were unique and unambiguous on each trial, so any sensory uncertainty was entirely due to the noise inherent in sensory processing. Bias was small but consistent when participants had ample time to process the target location and prepare their movement toward it to when they had ample time to view the target before moving, and when the location of targets was known without uncertainty in two-step movement tasks. This suggests that there is a component of movement bias toward previous actions even when sensory uncertainty is all but eliminated. However, when there was pressure to respond quickly to the target presentation, reaching and saccade direction biases were dramatically increased. Crucially, the bias magnitude correlated with trial-by-trial reaction time (i.e. preparation time) within time constraint conditions, implying that movement planning monotonically evolved from the prior location toward the target location upon presentation. This suggests that part of the mechanism for movement bias towards the prior distribution of targets arises due to advanced preparation of the action that is expected to be required.

In the current study, not only was bias greater when movement preparation time was short, but the degree to which sensory uncertainty influenced movement bias was also more consistent across participants. Nine out of ten participants increased bias with each reduction in sensory precision, suggesting that the fundamental feature of Bayesian inference that likelihood information should be inversely weighted by its precision was upheld despite limited time for time-consuming cognitive processing. This suggests that sensory precision can influence motor planning and execution via lowlevel sensorimotor transformation processes, as would be expected given the apparent ubiquity of Bayesian behavior throughout the animal kingdom (Berkes et al., 2011, Dekleva et al., 2016, Fischer and Pena, 2011). If we combine this finding with previous conclusions that movement bias under time pressure results from advanced preparation of the next expected action, we come to the conclusion that sensory uncertainty is integrated with movement preparation according to the principles of Bayesian inference.

The failure of participants in the long preparation group of the current study to show behaviour consistent with Bayesian principles suggest that the effects of sensory uncertainty are reduced with additional time for neural processing. This might occur because neural processing is better able to extract information from noisy sensory data with additional processing time, or merely because the effects of anticipatory motor preparation toward the prior have resolved by the time uncertain target information is transformed into a motor plan under long preparation time conditions. These possibilities remain speculative, however, because the defining feature of the behaviour shown by participants in the long-preparation group was the variability of reaching bias for different levels of sensory uncertainty.This variability stands in contrast to the orderly effect of sensory uncertainty on their basic motor variability. The standard deviation of movement directions systematically increased for with increasing sensory uncertainty for all subjects and was indistinguishable from that of the short preparation group.

Thus, the failure to display Bayesian behaviour was not because participants ignored the sensory uncertainty inherent in the target cue markers. Moreover, some participants did exhibit aiming biases that increased with increasing sensory uncertainty, which is consistent with the findings of Dekleva et al. (2016). We suspect that this idiosyncratic behaviour occurs because humans (and perhaps, occasionally, monkeys) apply sub-optimal cognitive strategies to anticipate or predict the likely target location from uncertain target information when they have ample time. There is a considerable literature illustrating that humans fail to behave rationally in decision-making tasks, for a variety of reasons (e.g. De Martino et al., 2006, Dayan and Niv, 2008, Rangel et al., 2008, Tom et al., 2007).Similarly, it would appear that providing enough time for people to make a conscious decision about how to integrate a novel form of noisy information about target location leads to suboptimal behavior. It remains to be seen whether people would eventually learn to apply Bayesian principles to their reaching behaviour with sufficient exposure to the task, and/or with sufficient incentive to reward good performance, but the results of Dekleva et al. (2016) suggest that this would be likely, given that their monkeys frequently exhibited Bayesian behaviour when repeatedly exposed to the task and provided juice rewards for target acquisition.

In summary, the current study showed that participants relied on prior target location information when target precision was reduced under time pressure, but displayed idiosyncratic behaviour when presented with amply time to process target information. This suggests that integration of target uncertainty information according to Bayesian principles is an inherent component of low-level sensorimotor transformation processes, but that statistically optimal behaviour can be undone by irrational cognitive processes if people are given enough (rope) time to apply them.

